# Photoperiod-dependent expression of MicroRNA in *Drosophila*

**DOI:** 10.1101/464180

**Authors:** Mirko Pegoraro, Eran Tauber

## Abstract

Like many other insects in temperate regions, *Drosophila melanogaster* exploits the photoperiod shortening that occurs during the autumn as an important cue to trigger a seasonal response. Flies survives the winter by entering a state of reproductive arrest (diapause), which drives relocation of resources from reproduction to survival. Here, we profiled the expression of microRNA (miRNA) in long and short photoperiods and identified seven differentially expressed miRNAs (*dme-mir-2b*, *dme-mir-11*, *dme-mir-34*, *dme-mir-274*, *dme-mir-184*, *dme-mir-184\** and *dme-mir-285*). Misexpression of *dme-mir-2b*, *dme-mir-184* and *dme-mir-274* in pigment-dispersing factor-expressing neurons largely disrupted the normal photoperiodic response, suggesting that these miRNAs play functional roles in photoperiodic timing. We also analyzed the targets of photoperiodic miRNA by both computational predication and by Argonaute-1- mediated immunoprecipitation of long- and short-day RNA samples. Together with global transcriptome profiling, our results expand existing data on other *Drosophila* species, identifying genes and pathways that are differentially regulated in different photoperiods and reproductive status. Our data suggest that post-transcriptional regulation by miRNA is an important facet of photoperiodic timing.

## Introduction

Adaptation to seasonal fluctuations in the environment drives some of the most incredible changes in animal behavior and physiology (Schwartz & Andrews 2013). Examples of common strategies for escaping unfavorable seasons include bird migration, insect diapause and mammalian hibernation. The accurate anticipation of the incoming season is vital for development, reproduction and fitness, and requires a seasonal timer (Bradshaw & Holzapfel 2010; Schiesari *et al.* 2011) that is often based on photoperiod measurement. The molecular details of this photoperiodic timer are still elusive, despite decades of intense research (Bradshaw & Holzapfel 2010;Saunders *et al.* 2004). Recently, however, significant progress has been made (Ojima *et al.* 2018; Andreatta *et al.* 2018; Anduaga *et al.* 2018; Liao *et al.* 2017).

In *Drosophila melanogaster*, the seasonal response is triggered by the short autumnal photoperiod and decreasing temperatures (Saunders *et al.* 1989). The response is manifested as an arrest of ovarian maturation in a pre-vitellogenic state that is characteristic of a reproductive diapause. The photoperiodic response in *D. melanogaster* is rather shallow, strain-specific and is largely masked by the temperature cycles (Anduaga *et al.* 2018). A recent study (Nagy *et al.* 2018) suggested that the photoperiodic response of flies in the laboratory could be greatly enhanced by adopting a semi-natural lighting regime, as opposed to the rectangular light-dark cycles which is commonly used. The photoperiodic response of *Drosophila* is also increased when the flies are starved (Ojima *et al.* 2018).

The mechanism of photoperiodic timing is still obscure, although recent evidence points to the rather old internal coincident model suggested by Pittendrigh (1972). In this model, the phase relationship between two circadian oscillators that are respectively driven by scotophase (night) and photophase (light) is used to encode the photoperiod. The circadian neuronal network in *Drosophila* consists of two distinct groups of clock neurons, each oscillating at a different phase (M-cells and E-cells) (Liang *et al.* 2016), which could also serve as a photoperiodic timer. M-cells produce pigment-dispersing factor (PDF) that is used to synchronize E-cells (via the PDF receptor) (Liang *et al.* 2016), while projections from M-cells and from insulin-producing cells (IPCs), which are essential for diapause activation, converge in the dorsal protocerebrum (Yasuyama & Meinertzhagen 2010). Remarkably, the loss of PDF in *Pdf^01^* mutants is sufficient to abolish the photoperiodic response (Ojima *et al.* 2018).

Activation of the diapause program in *D. melanogaster* involves insulin-like peptides secreted from the IPCs in the mid-brain, stimulating the *Corpora allatum* (CA, a subunit of the ring gland, a tripartite endocrine organ) to produce juvenile hormone (JH). JH promotes the synthesis of yolk proteins in the fat bodies (Q. Wu & Brown 2006). In the presence of JH, the ovaries are induced to produce ecdysteroid (Edc), which promotes yolk protein upload and vitellogenesis. The shutdown of JH and Edc determines the arrest of ovarian maturation in a pre-vitellogenic state that is characteristic of *D. melanogaster* diapause (Richard *et al.* 2001; Riddiford 2008).

MicroRNAs (miRNAs) are short non-coding RNAs which bind to the target 3’ untranslated region (UTR) of mRNA molecules and regulate expression of the encoded gene at the translational level (Gebert & MacRae 2018). Active and mature miRNAs are 17–24 base pair-long single-stranded RNA molecules that are expressed in eukaryotic cells and known to affect the translation or stability of target messenger RNAs (mRNA). MiRNAs are generated by cleavage of their pri-mRNA precursors by the endoribonuclease Drosha, followed by cleavage by Dicer in the cytoplasm. The miRNA is loaded into the RISC complex that contain Argonaute-1 (AGO-1), and then binds to target transcripts, particularly their 3’UTR regions. Binding of the miRNA to target region leads to either cleavage of the target transcript or inhibition of their translation.

Post-transcriptional regulation by miRNA has been implicated in a broad range of processes, including the circadian system (Xue & Zhang 2018). Recent studies in plants revealed that miRNAs are also involved in the photoperiodic timing of flowering (Jung *et al.* 2007; L.Wu *et al.* 2013), although no similar role in animal photoperiodism has been reported to date. Here, we report a global expression profiling of RNA extracted from female fly heads exposed to different photoperiods and diapause-inducing conditions and identify differentially expressed genes (DEGs) and miRNA (DEM) and demonstrate their functional role in photoperiodism.

## Methods

### Flies maintenance and samples collection

Canton-S flies were maintained in 200 ml glass bottles containing fly food (4.6% sugar, 4.6% brewer’s yeast, 1.25 % agar, 0.2 % methyl 4-hydroxybenzoate) at 25°C in a 12 h light-12 h dark cycle (LD 12:12). Flies were collected in a 6 h post-eclosion window and maintained at 12.5±0.2°C for long (LD 16:8) and short (LD 8:16) photoperiods (for temperature trace, see Fig S1), hereafter referred to as long day and short day, respectively. Twelve days later, female heads were collected in dry ice 4-6 h after light-on and stored at −80°C in TRIzol reagent (Invitrogen). Each sample was tested for the level of diapause based on previously described stages of ovary development (King 1970). A fly was considered to be in reproductive arrest if its most advanced oocyte was pre-vitellogenic (i.e., prior to stage 8). For over-expressing miRNA in *Pdf* neurons, we crossed UAS-miRNA flies (Bloomington stock number: 41128, 41172, 41174) with PDF-Gal4 flies (Bloomington stock number: 6900). As control experiments, both the UAS-miRNAs and PDF-Gal4 strains were crossed with flies presenting a *yellow, white* genetic background. The levels of diapause were recorded in these samples as described above.

### MiRNA and gene expression profiling

Total RNA was isolated using TRIzol reagent (Invitrogen). The miRNA fraction was isolated from each sample using the pureLink miRNA isolation kit (Invitrogen). Subsequently, the NCode multi-species miRNA microarray v2 (Invitrogen) was hybridized with fluorescently labeled miRNAs from flies experiencing both long and short days. The NCode multi-species miRNA microarray v2 alone was used as a positive control. The experiment was repeated 4 times. After hybridization, chip images were scanned using GenePix 4.1 software. Signal intensity was analysed using the Limma package of R software (Ritchie *et al.* 2015). Possible target genes for differentially expressed miRNAs were identified using Target scan (Ruby *et al.* 2007) and Pictar (Grun *et al.* 2005). DAVID software (Huang da *et al.* 2009) was used to search for enriched biological functions amongst the identified targets.

### AGO-1 immunoprecipitation and RNA sequencing

AGO-1 immunoprecipitation was carried out using a previously described protocol (Kadener *et al.* 2009) with minor modifications (See Supplementary methods). RNA-seq library preparation and sequencing were carried out by Glasgow Polyomics (www.polyomics.gla.ac.uk) using the IlluminaNextseq500 platform. Two independent libraries (pair-end) for long day and one for short day (input and IP) flies were generated. Adapter sequences and poor-quality readings were trimmed from the RNA-seq data, and the quality of the resulting trimmed libraries was checked with fastQC (0.11.2) (Andrews S 2010).

## Results

### Differential gene expression

We compared gene expression in the heads of diapausing and non-diapausing females and identified 143 DEGs (p<0.01; Fig 1, Table S1). Only few biological functions were significantly enriched among these genes, including those encoding ahydrolase, peptidase M13 (Benjamini p<0.05) and signal peptide processing (Benjamini p =0.056; Fig S2). In the photoperiodic experiment, we identified 517 DEGs. Among the top hits were *Hsp70* proteins which were previously implicated in insect diapause (Rinehart *et al.* 2007). Other interesting genes identified were *vrille* (*vri*) ((Yuan *et al.* 2005; Martinek *et al.* 2001; Panda *et al.* 2002; Stanewsky 2002; Cyran *et al.* 2003), and *shaggy* (*sgg*)(Martinek *et al.* 2001), clock genes that may serve as a link between the circadian and photoperiodic clocks. Over-represented processes among the photoperiodic DEGs included ‘oxidoreductase’, ‘alternative splicing’, and ‘fatty acid and acyl-CoA metabolism’ (Fig. S2).

**Figure 1.**
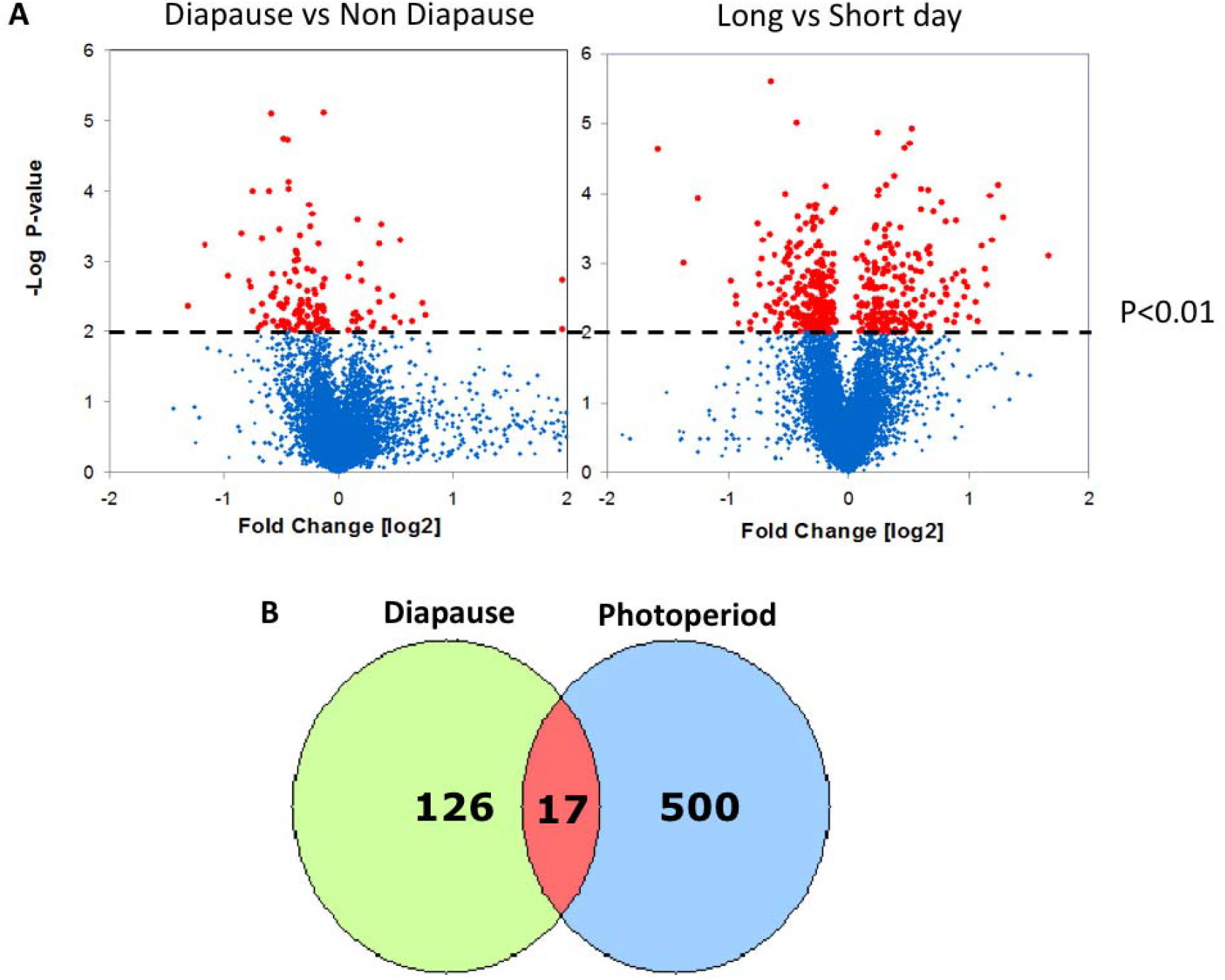
Expression profiles. **A.** Volcano plots showing differential expression in female heads (p<0.01, red). In the diapause experiment (left), RNA was extracted from female heads following 12 days in LD (14:10) conditions at 12±0.2ºC,with positive fold changes representing up-regulation in diapause. In the photoperiodic experiment (right), females were maintained at LD (16:8) and SD (8:16) conditions, with positive fold changes representing up-regulation in long day readings. **B.** Overlapping Venn diagrams of photoperiodic and diapause DEGs.

The intersect of the two lists reveals 17 transcripts that may serve as candidate loci for linking the photoperiodic timer with diapause induction (Table S2). These included genes encoding yolk proteins 2-3, deoxyribonuclease II, CG3829, CG1CG1 G3829, CG10516, CR17875 (heme-binding) and Obp99a. The gene encoding another odorant-binding protein in the photoperiodic list (Obp99b; Table S1) is known to be regulated by JH (Yamamoto *et al.* 2013; Andreatta *et al.* 2018).

### miRNA Expression

The expression of miRNA in fly heads exposed to ether long or short photoperiods was carried out using microarrays (Fig. 2). We detected seven differentially expressed miRNA (DEMs) that included *dme-mir-2b*, *dme-mir-11*, *dme-mir-34*, *dme-mir-274*, *dme-mir-184*, *dme-mir-285* (all at p < 0.01) and *dme-mir-184\** (at p = 0.01). We have also measured the expression of *dme-mir-274* and *dme-mir-184* by qPCR. Although the fold-change of both miRNAs correlated with the results of the array experiment, they were not statistically significant (note that for each miRNA, the t test power (1- β) was less than 10% to detect the observed difference).

To test for biological function enrichment among the miRNA targets, we analyzed the gene ontologies of the top 20 genes targeted by the DEMs (Table 1, Fig.2). We found significant enrichment of processes such as ‘septate junction’, ‘membrane activity’ and ‘negative regulation of transcription’. A number of genes were also targeted by more than one miRNA (Table 1), although their function in diapause or a photoperiodic response has yet to be elucidated. Overall, these results suggested that miRNAs may play an important regulatory role in the fly photoperiodic response.

**Table 1.**
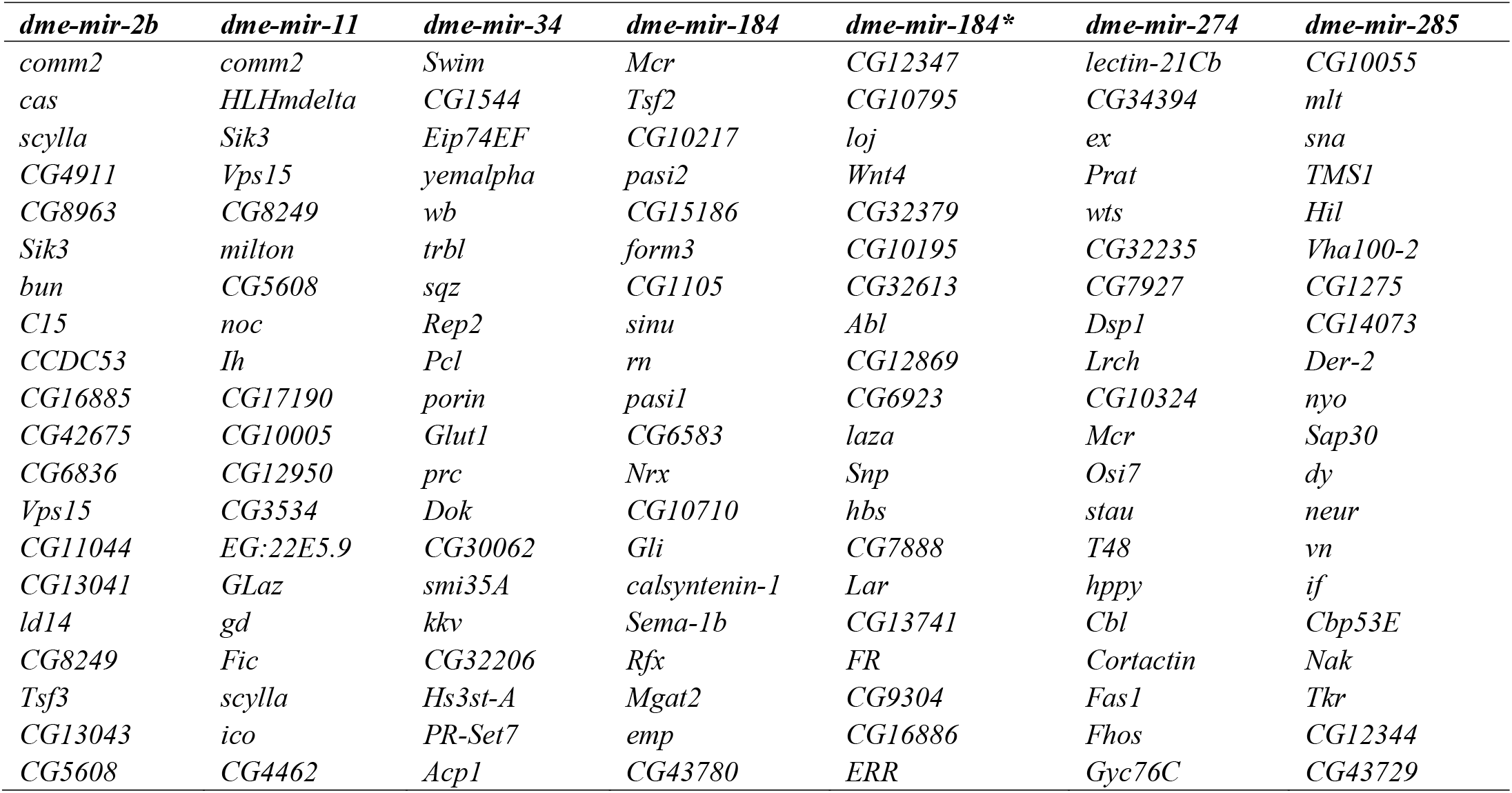
Top 20 predicted targets of the differentially expressed miRNAs.

**Figure 2.**
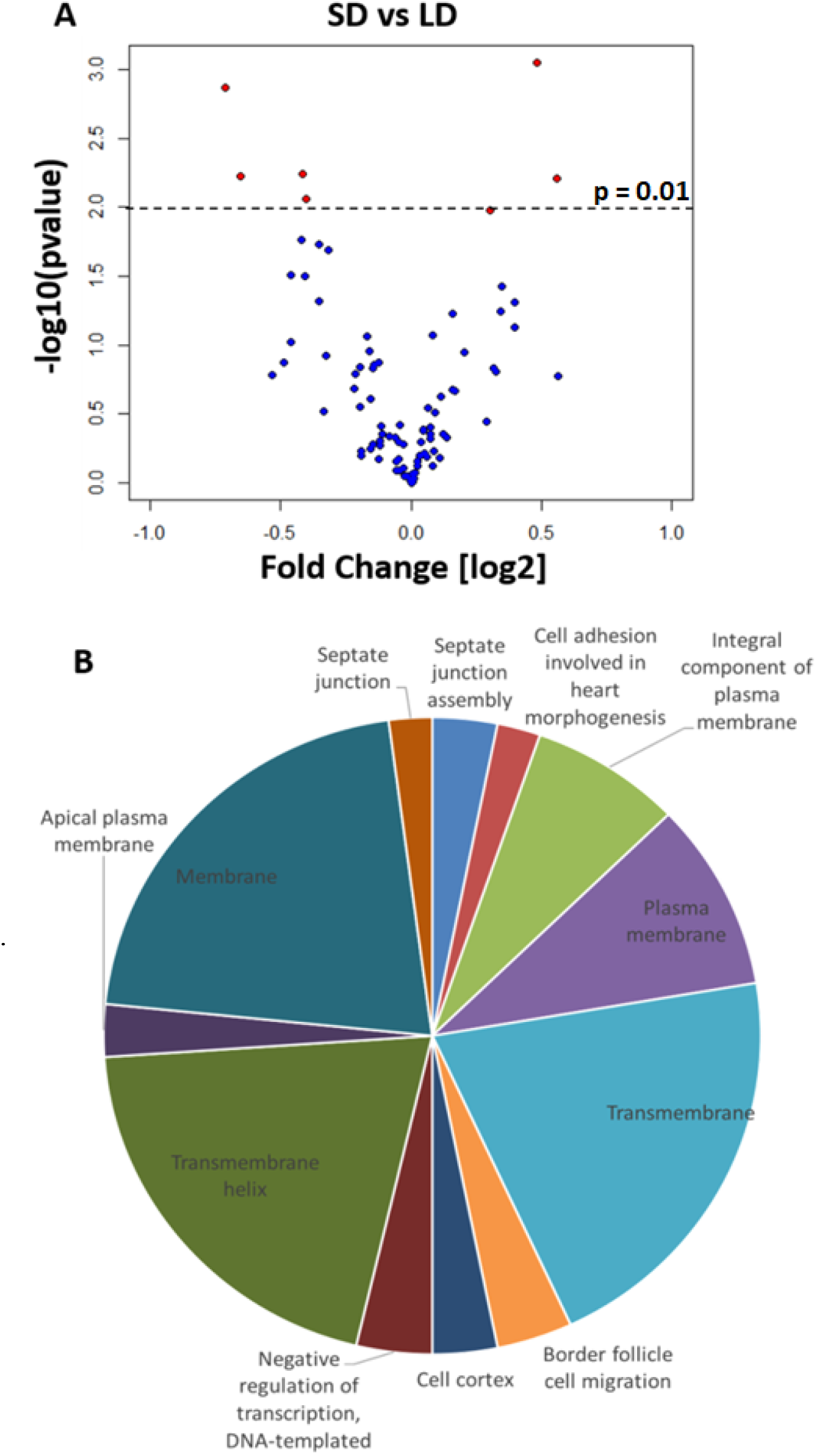
Differences in miRNA expression. **A.** Volcano plot showing differentially expressed miRNA in female heads (p<0.01, red; *dme*-*mir184\** p =0.0104) exposed to long (LD) and short (SD) day conditions. Positive fold changes indicate up-regulation in short day readings. **B.** Pie chart showing enriched biological functions (DAVID, Benjamini p<0.05) of the 20 most probable targets for each DEM. The size of the pie section is proportional to the number of genes for that biological function. SD: short day. LD: long day.

### Network Analysis

To identify the pathways associated with the photoperiodic response, we generated a network of protein-protein interactions (Shannon *et al.* 2003; Murali *et al.* 2011) combining the DEG lists (Diapause and photoperiodic) with the top 20 target genes of each DEM. The network analysis identified functions not immediately evident from the microarray results. Biological functions, such as ‘regulation of alternative mRNA splicing’, ‘chromatin silencing’, ‘regulation of Notch signaling pathway’, ‘courtship behaviour’ and ‘phototransduction’, were shared by the three microarray experiments (Table S3). Some biological functions were unique for a particular array. The photoperiodic network consisted of biological functions such as ‘rRNA methylation’, ‘histone phosphorylation’, ‘gene silencing by miRNA’ and ‘regulation of circadian sleep/wake’ (Table S3). ‘Entrainment of the circadian clock’, ‘regulation of circadian rhythm’, ‘response to heat’, ‘visual behavior’ and ‘JAK-STAT cascade’ were represented in both the miRNA and photoperiodic networks (Table S3). The diapause network contained just one unique biological function (‘fat body development’) but shared many biological function with both the photoperiodic network (‘tRNA methylation’, ‘regulation of translational initiation’ and ‘phospholipid biosynthetic process’) and the miRNA network (‘ecdysone receptor-mediated signaling pathway’, ‘positive regulation of hormone secretion’ and ‘rhodopsin-mediated signaling pathway’). A unique biological function of the miRNA network was ‘mitochondrial electron transport’ (Table S3). We generated another network intersecting the photoperiodic, diapause and miRNA datasets, which revealed two main pathways (Fig 3). The central node of the largest graph was cdc2-related-kinase, encoded by a diapause DEG. In the second graph, a photoperiodic DEG, a diapause DEG and a DEM all interacted with the central node (the C-terminal-binding protein gene), suggesting that a minute change in expression of just a few genes can have a cascading effect on a larger gene network. The biological functions represented in the larger interaction network were ‘fatty acid biosynthetic process’, ‘positive regulation of hormone secretion’, ‘regulation of transcription’, ‘regulation of circadian sleep/wake cycle’, ‘positive regulation of apoptosis’, ‘ecdysone receptor-mediated signaling’, ‘determination of adult lifespan’ and ‘nuclear mRNA splicing’ (Fig 3). The biological functions in the second network were ‘regulation of transcription’, ‘olfactory behavior’ and ‘Notch signaling pathway’ (Fig 3).

**Figure 3.**
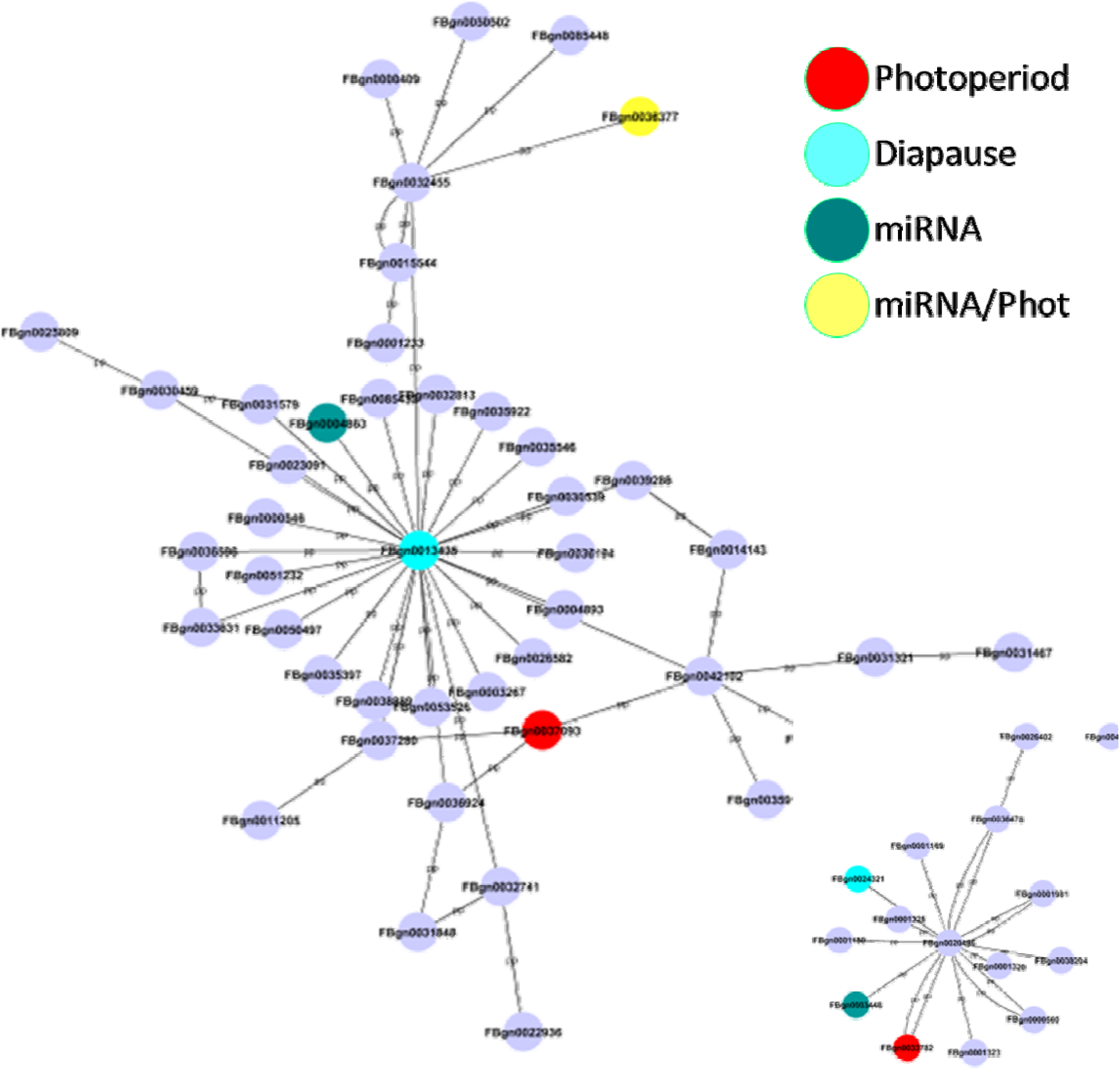
Intersection of Yeast Two-Hybrid interaction networks. The intersection of the three Yeast Two-Hybrid interaction networks (Diapause, Photoperiodic and miRNA) resulted in two interaction pathways. The largest interaction pathway (top) has at its center at *cdc2-related-kinase*, a diapause DEG (Table S1). The second interaction pathway has at its center at the gene for the C-terminal-binding protein. Red nodes are photoperiodic DEGs, light blue nodes are diapause DEGs, green nodes are DEMs targets and yellow nodes are both DEMs targets and photoperiodic DEGs.

### miRNA misexpression

To explore the functional roles of DEMs in the photoperiodic response, we induced the over-expression of *dme-mir-2b*, *dme-mir-184* and *dme-mir-274* using UAS transgenes driven either by strains carrying *Act*- Gal4 (ubiquitous expression) or *pdf*-Gal4 (in *pdf-*expressing neurons), and tested for any changes in photoperiodic diapause. We compared changes in photoperiodic responses between control (i.e., flies carrying a single transgene) and over-expressing (i.e., flies carrying both transgenes) lines by fitting a generalized ANOVA linear model. A significant photoperiod × genotype interaction was considered as being a significant effect of such misexpression. The over-expression induced by *Act*-Gal4 resulted in almost total lethality (data not shown), while over-expression in PDF-expressing neurons resulted in a loss of the photoperiodic response (Fig 4). Misexpression of *mir274* abolished the difference between long- and short-day flies (compared to UAS-*mir274* control, χ^2^ _1,16_ =5.51, p<0.05 and *pdf*-Gal4 χ^2^ _1,16_ =11.28, p<0.001), indicating a substantial reduction in short day diapause compared to controls (Fig 4). Over-expression of *dme-mir-2b* resulted in a similar outcome (*Pdf*-Gal4>mir2b vs. UAS-mir2b, χ^2^_1,18_= 12.26, p<0.001; *Pdf*-Gal4>mir2b vs. *Pdf*-Gal4, χ^2^_1,16_=17.05, p<0.001), while in the case of *mir184*, the short day reduction was accompanied by an increased percentage of long day diapause (Fig 4; *Pdf*- Gal4>mir184 vs. UAS-*mir184*, χ^2^_1,19_=11.58, p<0.001; and vs. *Pdf*-Gal4>Yw, χ^2^_1,18_=25.52, p<0.001).

**Figure 4.**
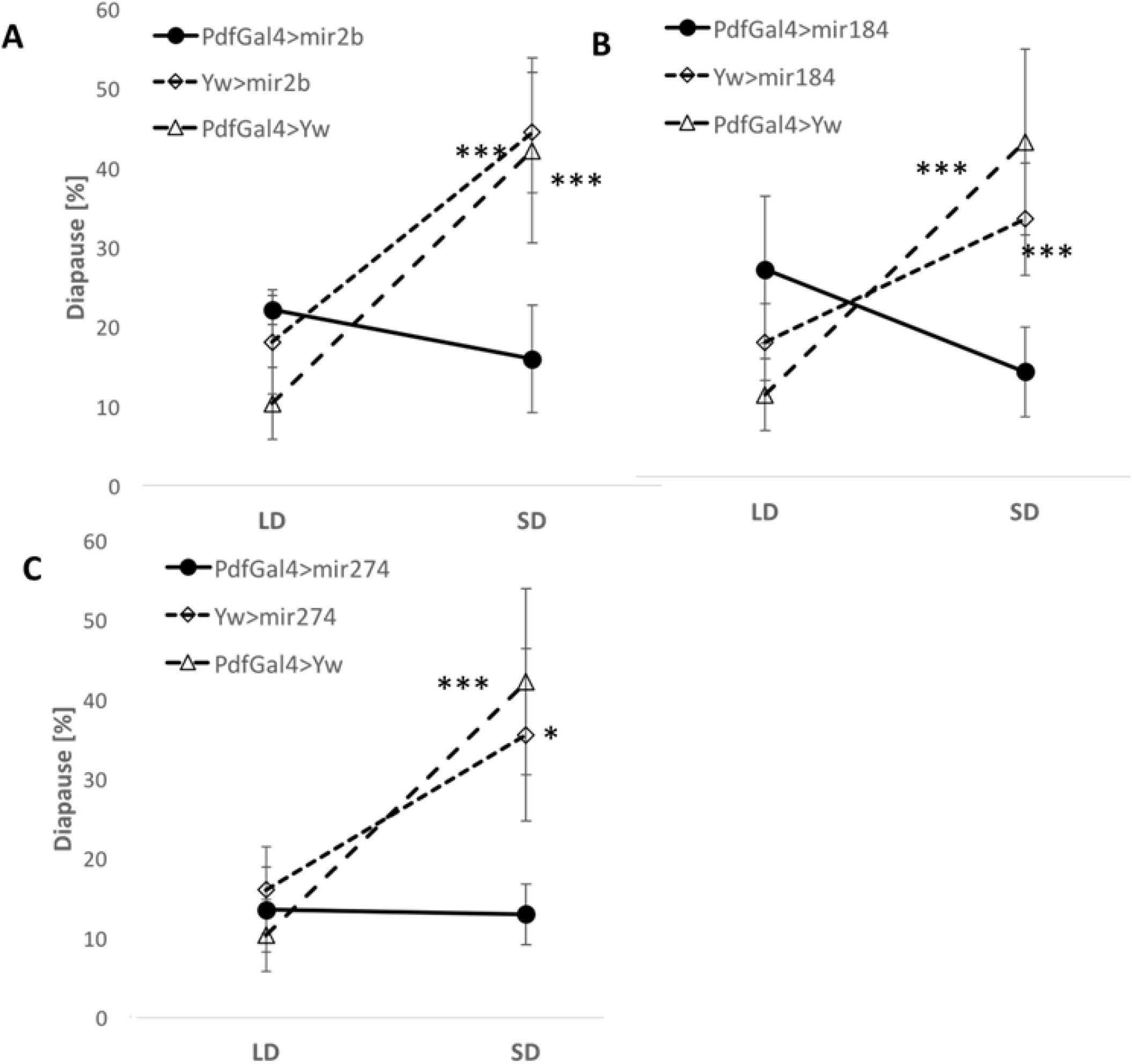
miRNA over-expression causes a loss of photoperiodic response. Average diapause [%] in flies over-expressing (**A**) *dme-mir-2b*, (**B**) *dme-mir-184* and (**C**) *dme-mir-274*, in PDF-expressing cells. Error bars represent SEM. Significant Phot:Gen interactions [p<0.001(***), p<0.05(*)] indicate a different photoperiodic behavior between over-expressing lines and controls.

### AGO-1 immunoprecipitation

We used AGO-1 immunoprecipitation, followed by sequencing of the associated RNA, and tested for transcripts enriched in either long- or short-day samples. We found 643 genes whose transcripts were significantly enriched in long day immunoprecipitates, and 68 genes that were enriched in short day immunoprecipitates (q < 0.05, Table S3). Among the long-day genes were 62 over-represented biological functions (FDR < 0.05; Fig S3) including ‘alternative splicing’, ‘regulation of transcription’, ‘zinc-finger’, ‘transmembrane activity’, ‘septate junction’ and ‘eye development’. In contrast, in the short day gene list, ‘ether lipid metabolism’ was the only term deemed marginally significant (FDR = 0.058; Fig S3). A number of targeted genes were particularly interesting. The long day list included genes like *AGO1*, *ecdysone receptor* (*EcR*), and circadian clock genes, such as *clock* (*Clk*), *pdp1* and *vri*. Genes of the insulin pathway were also enriched in the long day preparation (i.e., *InR* and *Ilp6*). In addition, some predicted DEM targets were present in the long day list (i.e., *comm2*, *Tsf2*, *Hs3st*-A, *sinu*, *CG43780*, Nrx-IV, *Pcl* and *trbl*) and in the short day list (*sinu*). Interestingly, insulin-like receptor (*InR*) is one of the 51 enriched genes common to both the short and long day immunoprecipitate lists.

## Discussion

This study was aimed at testing the role of miRNA in the photoperiodic response of *Drosophila*. We identified seven miRNA that were differentially expressed between long and short photoperiods (i.e., *dme-mir-2b, dme-mir-11, dme-mir-34, dme-mir-274, dme-mir-184, dme-mir-184*, dme-mir-285*). The roles and functions of some of these miRNAs in *Drosophila* are not clear yet (such as *dme-mir-2b, dme-mir-274, dme-mir-285*). However, the biological processes affected by *dme-mir-11, dme-mir-34 and dme-mir-184* show an intriguing link to photoperiodism and diapause. *dme-mir-11*, for example, was implicated in regulating apoptosis during embryonic development (Ge *et al.* 2012) and in regulating pupal size during metamorphosis (Y.Li *et al.* 2017). Interestingly, *dme-mir-34* plays an important role in early neuronal differentiation and ageing (Soni *et al.* 2013; Liu *et al.* 2012), as well as in ecdysone signaling (Xiong *et al.* 2016). *dme-mir-184* is involved in oogenesis, development and ovary morphogenesis (Iovino *et al.* 2009; P.Li *et al.* 2011; H.Yang *et al.* 2016), while the daily expression of this miRNA shows circadian oscillation (M.Yang *et al.* 2008). Over-expressing *dme-mir-2b, dme-mir-184* and *dme-mir-274* using *Act*- Gal4, a driver inducing spatially and temporally broad expression, resulted in almost total lethality, suggesting central roles for these three miRNAs during embryogenesis/development. Remarkably, over-expression of these miRNAs in PDF-expressing cells resulted in a loss of photoperiodic response, albeit in different ways. Our microarray and real-time PCR experiments suggested that both *dme*-*mir-2b* and *dme-mir-274* are expressed at higher levels in long days than in short days. Consistent with this observation, over-expression resulted in a substantial reduction of diapause in short days, when there is normally a lower level of these miRNA (as compared to long days). *dme-mir184* is highly expressed in short days, with its over-expression resulting in a reduction of short day diapause, accompanied by an increase of long day diapause and a reversal of the normal photoperiodic response. These data suggest a possible causative role of these miRNAs in the photoperiodic response.

Recently, dme-*mir-210* was implicated in mediating clock function and light perception, possibly affecting the arborization of the ventral lateral neurons (LNv, pdf+ cells) (Cusumano *et al.* 2018; Ojima *et al.* 2018). It was suggested (Ojima *et al.* 2018) that light information reaches the IPCs via PDF-expressing cells. As such, it is possible that controlling miRNA expression in these cells is either integral part of the photoperiodic time measurement or miRNA drives a cascading affecting its expression. Our AGO-1 immunoprecipiation experiment revealed transcripts that were differentially enriched in the immunoprecipitates, some of which were predicted targets of DEMs that we identified (Table 5). In addition, the miRNA microarray experiment suggests that there are more miRNAs over-expressed in long day than in short day flies (Fig 3). Consistently with this observation, the long day AGO-1 immunoprecipitate contained almost 10 times more enriched genes than did the short day immunoprecipitate (643 vs. 68; Table S3). Interestingly, ‘septate junction’ and ‘membrane activity’ functions were enriched both amongst the DEMs targets and in the long day AGO-1 immunoprecipitated genes.

In addition, we identified 143 genes differentially expressed between diapausing and non-diapausing females (Fig 1, Table S1). Lirakis *et al.* (Lirakis *et al.* 2018) suggested re-define diapausing as a stress response to cold temperatures. They suggested that diapause-scoring methods, like the one applied here, may underestimate the true degree of diapause. In fact, they showed that oogenesis was blocked at either a previtellogenic or a very early vitellogenic stage when flies were exposed to dormancy-inducing conditions. This might be the reason underlying the relatively modest number of DEGs seen, as compared to the photoperiodic experiment. Only few biological functions were enriched in our diapause DEGs (Fig 2), with some genes being clearly related to diapause (e.g. *yp2-3*, *Egfr*). In the photoperiodic experiment, the number of DEGs was substantially higher (517; Fig 1, Table S1). Over-represented functions in this list included ‘oxidoreduction’, ‘acyl-CoA metabolism’ and in particular, ‘alternative splicing’ (Fig S2). ‘Alternative splicing’ and ‘splice variants’ were also enriched functions among the long day AGO-1 immunoprecipitate transcripts. Alternative splicing of circadian clock genes has been previously implicated in the response to seasonal changes, be they photoperiods or temperatures (Glaser & Stanewsky 2005;Montelli *et al.* 2015; Collins *et al.* 2004; Chen *et al.* 2007). Our results expand these data, suggesting that alternative splicing may be a more general strategy to allows a plastic response to environmental changes in *Drosophila*.

Whether the circadian clock is required for photoperiodic timing in flies is not yet clear, although recent evidence suggests a photoperiodic role for clock PDF-expressing neurons (Ojima *et al.* 2018). Our list of photoperiod DEGs included *vri* and *Shaggy* (*Sgg*) that may serve as a link between the photoperiod and circadian clocks (Yuan *et al.* 2005; Martinek *et al.* 2001; Panda *et al.* 2002; Stanewsky 2002; Cyran *et al.* 2003; Stanewsky 2003).

Two clock genes (*Clk* and *vri*) were also among the miRNA targets (long day) that we identified by the AGO-1 immunoprecipitation. In a screen for diapause genes in *D. montana*, *cpo* and the circadian clock genes *vri* and *per* were seen to be differentially expressed (Salminen *et al.* 2015). *per* and *cpo* were associated with the diapause initiation phase, while *vri* was associated with the diapause maintenance phase. In agreement with Salmien *et al*. (Salminen *et al.* 2015), our photoperiodic DEGs included genes encoding the myosin heavy chain (Mhc), pyrroline 5-carboxylate reductase (P5cr2) and Hsp70s. Interestingly, our photoperiodic DEGs also included a number of odorant-binding protein genes (i.e., *Obp56g*, *Obp99a*, *Obp99b*, *Obp83g* and *Obp49a*). Similar genes have been previously associated with diapause in *D. montana* (Kankare *et al.* 2016). *Obp99b* is repressed by JH, which is produced by the CA (Yamamoto *et al.* 2013) and was found to be up-regulated in diapause (Andreatta *et al.* 2018).

The network analysis presented here revealed additional pathways that are associated with both diapause and the photoperiodic response. A unique function that was enriched among diapause DEGs was ‘fat body development’. Indeed, the fat bodies are responsible for the production of yolk proteins (YP) that are fundamental components of the developing oocyte. During diapause, the production of ILPs is repressed, resulting in a repression of JH and Ecd expression, which in turn inhibits the synthesis of YP in the fat bodies (Schiesari *et al.* 2011). Other pathways that were shared between the miRNA and photoperiodic networks fit very well with the expected diapause-hormonal axis. These pathways include ‘ecdysone receptor-mediated signaling’, ‘positive regulation of hormone secretion’ and ‘phospholipid biosynthetic process’. The pathway ‘rhodopsin mediated signaling’ was shared by both the diapause and miRNA networks, and may constitute a possible link between light perception (compound eyes, ocelli) and the IPCs responsible for ILP production. *EcR* and other genes of the insulin pathway (i.e., *InR*, *Ilp6*; Table S2) were indeed enriched in long day flies in our AGO immunoprecipitation experiments. The photoperiodic network included unique biological processes such as ‘rRNA methylation’, ‘histone phosphorylation’, ‘regulation of circadian sleep/wake’ and ‘gene silencing by miRNA’ (Table 3). The other functions associated with this network underscore the significance of mechanisms that regulate transcription (e.g. alternative splicing histone modifications, miRNA-dependent silencing) to the photoperiodic response. Functions shared by the diapause and the photoperiodic networks (i.e., ‘tRNA methylation’ and ‘regulation of translational initiation’) reinforce this notion. Interestingly, biological functions shared by both the photoperiodic and miRNA networks (i.e., ‘entrainment of circadian clock’, ‘regulation of circadian rhythm’, ‘response to heat’ and ‘visual behavior’) (Table 3) were also associated with the diapause response in *D. virillis* (Salminen *et al.* 2015). Intersecting all the networks produced two interaction hubs of genes (Fig 3), represented in a number of important functions, including ‘fatty acid biosynthetic process’, ‘positive regulation of hormone secretion’, ‘regulation of transcription’, ‘regulation of circadian sleep/wake cycle’, ‘positive regulation of apoptosis’, ‘ecdysone receptor-mediated signaling’, ‘determination of adult lifespan’, ‘nuclear mRNA splicing’, ‘olfactory behavior’ and ‘Notch signaling’. Finally, biological functions, such as ‘acid biosynthetic’, ‘acyl-CoA metabolism’, ‘oxido-reduction’, that we identified here were associated with diapause in previous studies in other *Drosophila* species (Salminen *et al.* 2015; Kankare *et al.* 2016).

